# Extending protein interaction networks using proteoforms and small molecules

**DOI:** 10.1101/2022.09.06.506730

**Authors:** Luis Francisco Hernández Sánchez, Bram Burger, Rodrigo Alexander Castro Campos, Stefan Johansson, Pål Rasmus Njølstad, Harald Barsnes, Marc Vaudel

## Abstract

Biological network analysis is used to interpret modern high-throughput biomedical data sets in terms of biological functions and pathways. However, the results greatly depend on the topological characteristics of the underlying network, commonly composed of nodes representing genes or proteins that are connected by edges when interacting. In this study, we build biological networks accounting for small molecules, protein isoforms and post-translational modifications. We highlight how these change the global structure of the network and how the connectedness of pathway-based networks is altered. Our findings highlight the importance of carefully crafting the networks for network analysis to better represent the reality of biological systems.

## BACKGROUND

Biological networks are a promising way to interpret modern biomedical data at scale^1^. They allow the study of molecular patterns at both local and global scale, and hence provide a systemic view on molecular processes. For example, by modeling the interactome — the entire collection of biological interactions — Menche *et al*. identified disease modules, and studied their topological properties and pairwise relationships^2^. An overlap between disease modules would then indicate a functional relationship, hinting at shared mechanisms and possible common drug targets.

The fundamental building blocks of a biological network are the interactions between biological entities, with the entities themselves represented by nodes and their interactions by connections^3^. The entire collection of interactions in a biological system is called the interactome. The main participants of the interactome are proteins, represented in biological networks by the name of the gene encoding them. A relationship between proteins can be inferred from multiple sources: text mining, co-expression, physical interaction, or from literature knowledge on the functions of proteins^4^. Such networks have proved to be very useful for understanding biological mechanisms^5–7^. For example, gene network approaches have been used for analyzing functions of genes associated to different types of cancer^8,9^.

Based on a given interactome, network analyses attempt to extract knowledge concerning specific sets of proteins. For example, *guilty by association* procedures assume that proteins colocalizing in the network are functionally related^8^. Similarly, diffusion models estimate the effects of gene alterations towards their neighborhood^10^. By design, such network analysis methods rely heavily on network structural properties such as the number of neighbors per node or the number of connections between groups of nodes^1^. It is then vital to carefully choose what the nodes and connections represent, such that any inference from the network mirrors the reality of biological systems.

In practice, as a result of genetic variation, RNA splicing, and post-translational modification (PTM), a gene can yield many distinct forms of a protein, called proteoforms^11^. For most proteins, the different isoforms of a gene share less than 50 % of interactions^12^. For example, *Bcl-2* has two isoform products Bcl-xl and Bcl-xs resulting from alternative splicing. Bcl-xl, which contains the BH1 and BH2 domains, is responsible for programmed cell death, while Bcl-xs lacks both domains, therefore contributing to the opposite function^13^. One can legitimately anticipate an even higher specificity when including PTMs. However, this information is lost when creating biological networks using gene names as sole descriptor of the protein.

Another source of information lost in the construction of gene-centric networks is the role of small molecules, which play essential roles in biological systems, *e.g*., metabolites participating as reactants, catalyzer, or inhibitor of reactions. For example, adenosine triphosphate (ATP) and guanosine triphosphate (GTP) are essential metabolites needed as energy sources. ATP hydrolysis provides the energy for protein transport in the mitochondria, for binding and releasing the newly synthesized polypeptide molecules from the *hsp70* chaperone proteins^14^.

Previously, we have demonstrated that it is possible to leverage the rich information contained in the Reactome pathway knowledgebase to refine the representation of biological networks by accounting for proteoform-specificity of biological reactions^15^. Here, we demonstrate how changing the type of node from gene to proteoform influences the structure of the obtained networks. In addition, we study how the inclusion of small molecules affects the representation of the network. Together, our results show that changing the representation biological networks can help refine the modeling of biological processes, but that the limited information of proteoform-specific interaction still impairs the application of such approaches at scale.

## RESULTS

### INCREASED SIZE OF THE INTERACTOME

A recent estimate for the human genome lists approximately 47,000 genes, of which approximately 19,000 are coding for proteins^16^. The estimated number protein products resulting from alternative splicing is around 70,000 isoforms^17^. The total number of functional proteoforms remains unknown, but estimates are in the millions depending on how proteoforms are defined^18^. Changing the representation of a network from a gene-centric to a proteoform-centric paradigm should therefore result in a network several orders of magnitude larger. Based on isoform and post-translational modification information from the Reactome knowledgebase v80 for *Homo Sapiens*, we can represent 14,246 distinct proteoforms participating in 13,806 reactions (see methods for details). These 14,246 proteoforms represent 11,074 proteins linking to 10,976 gene names, making 1.3 proteoforms per gene on average. We constructed a network based on all pathways in Reactome by connecting entities when they participate in the same reaction. Building the network based on proteoforms instead of genes yields 3,270 (+29.8 %) additional nodes and 224,207 (+61.2 %) additional connections. Thus, while the proteoform annotation provides enough information to substantially increase the size of the network, only few proteoforms are annotated functionally.

The genes with the highest number of proteoforms annotated are *UBC*, *H3C1*, and *H3C15*, with 55, 52, and 48 proteoforms respectively, participating in diverse pathways and located in multiple subcellular compartments (**Supplementary Table 1**). *UBC*, for example, has products mostly ubiquitinylated or with crosslinks between L-lysine residues and glycine at multiple locations of the sequence, generating a high number of proteoforms representing different combinations of post-translational modifications. *HLA-A* and *HLA-B* are also genes with high numbers of proteoforms, not due to splicing variants or PTMs, but because there are multiple protein accessions linked to them, 36 and 21, respectively. For these examples, the proteoform representation of biological interactions will be completely different compared to a gene-centric network.

Reactome contains also small molecules annotated as participants of human reactions. Extending the gene- and proteoform-centric networks with small molecules increases the number of nodes by 2,057, representing an increase of 18.7 % and 14.4 %, respectively (**Supplementary Table 2**). Adding small molecules creates 85,282 and 91,476 new connections, corresponding to an increase of 23.3 % and 15.5 % for the gene- and proteoform-centric networks, respectively. However, this creates situations where small molecules ubiquitous in biochemical reactions, like H_2_O or ATP, connect most of the network. To take the influence of small molecules into account without distorting the network globally, we introduced the possibility for small molecules to connect pathway participants within but not between reactions. The number of new connections then becomes 442,004 and 457,127, corresponding to an increase of 120.7 % and 77.4 %, for the gene- and proteoform-centric networks, respectively.

### INTERCONNECTED PROTEOFORMS ALTER THE DEGREE DISTRIBUTION

The connectivity of a node in a network is measured by the number of connections, also called the degree. Without accounting for small molecules, 143,255 (24.3 %) of the connections in the proteoform-centric network represent connections where proteoform-level information is available for both nodes, while 101,907 (17.3 %) and 345,253 (58.5 %) of the connections present no proteoform annotation for one or both nodes, respectively. Since proteoforms are specific forms of a protein^11^, changing a gene-centric network into a proteoform-centric representation can be seen as distributing the protein-protein interactions between new, more specific, nodes. Thus, intuitively, proteoform nodes are expected to a have a smaller degree than the gene that encodes them. As we previously described^15^, the majority of proteoforms indeed present a degree lower than their genes in the gene-centric network. This picture is however complicated by proteoform-proteoform interactions and, as detailed in **Figure 1** and **Supplementary Table 3**, at the scale of the entire network, the degree is increased when taking proteoforms into account.

**Figure 1.**
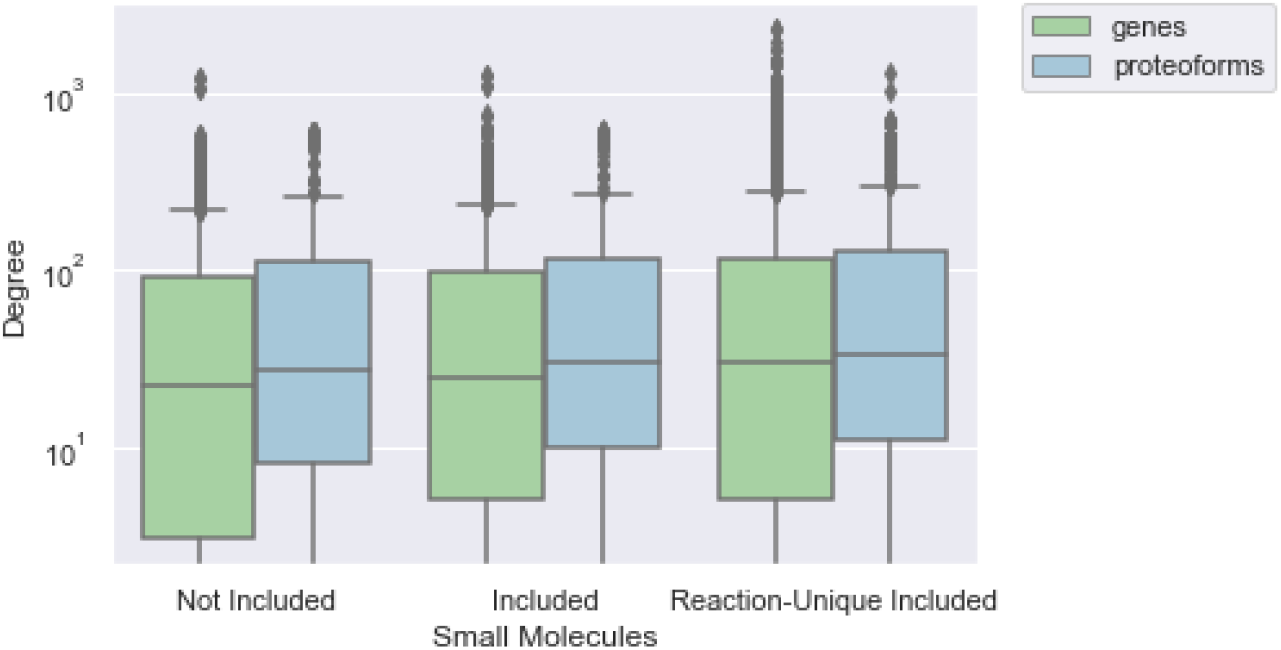
Node degree distribution in the different interactomes depending on how small molecules are considered. Left: only gene or proteoform nodes and not including small molecules as nodes. Center: small molecules included, one node for each. Right: “reaction-unique” small molecule nodes included, adding one separate node for each reaction where the small molecule participates (“reaction-unique”).

This is for example the case for collagen-related genes such as *COL7A1*, *COL3A1*, and *COL6A3*, which present a much higher degree in the proteoform-centric than in the gene-centric network: 606 vs. 121, 547 vs. 67, and 546 vs. 66, respectively. These collagen nodes are expanded to a wide variety of proteoforms as they become multiply modified by sequential reactions. For example, in the pathway *Collagen biosynthesis* a reaction converts *collagen lysines* to *5-hydroxylysines*, and diverse *COL7A1* gene products are input and output of the reaction. In a gene-centric network, this reaction is modeled as a single *COL7A1* gene node, while in the proteoform-centric network, the input nodes *COL7A1, 3×4Hyp-COL7A1*, and *3×4Hyp-3Hyp-COL7A1* are connected to the output nodes *5Hyl-COL7A1, 3×4Hyp-5Hyl-COL7A1*, and *3×4Hyp-3Hyp-5Hyl-COL7A1*, yielding nodes with higher detail of information but also with higher degree than in the gene-centric network. Other nodes that consequently have their degree increased do not necessarily have proteoform-level annotation, such as *PLOD3*, which has its degree increased by an order of magnitude (from 46 to 529), simply because it participates in reactions with multiple collagen gene products, therefore connecting to many proteoforms. The examples of genes with highest increase in degree between gene- and proteoform-centric networks are listed in **Supplementary Table 4**.

To evaluate the local vs. global effect of introducing proteoforms in the network, we evaluated the degree of nodes per pathway (**Supplementary Table 5)**. Then, the average degree per proteoform, 14.1, was slightly lower than per gene, 14.3 (−1.4 %). It therefore appears that the increase in degree observed for the whole network is not due to within-reaction or within-pathway connections, but rather between-pathway connections between proteoforms and other proteins. This highlights the importance of between-pathway connections and how picturing canonical pathways as separate entities distorts the reality of the interactome. We further evaluated whether the robustness of the network was altered by introducing proteoforms using a percolation analysis^2^. Both gene- and proteoform-level interactomes showed similar percolation curves, with a slightly better robustness for the proteoform network **Figure 2**.

**Figure 2.**
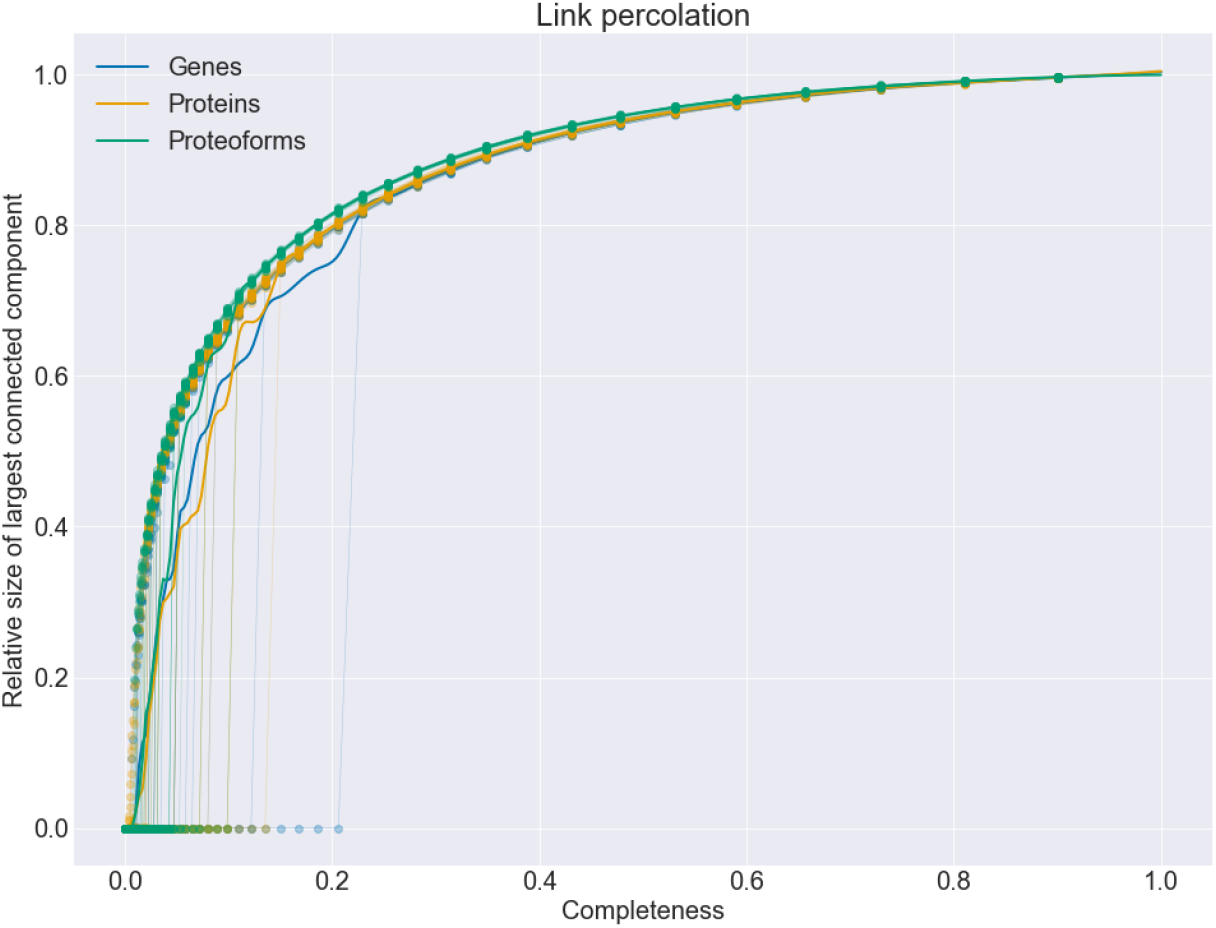
Approximations of link percolation curves for gene, protein, and proteoform interactome networks without small molecules. Relative size of the LCC (y axis) is the number of nodes with relatively to the original number of nodes in the complete network. Dots represent measures at each step of the percolation. Soft colored lines connect measures of each replicate, starting with completeness 1.0 (complete interactome) and iteratively removing connections until completeness is 0. Intense lines show average tendencies. Completeness (x axis) is the share of original connections kept after removing random connections.

As detailed in **Supplementary Table 3**, when extending the gene- and proteoform-centric networks with small molecules, the average degree of accessioned entity nodes increased from 66.7 to 72.9 (+9.2 %) and from 82.9 to 88.1 (+6.2 %), respectively. As previously introduced, the ubiquitousness of small molecules however produces hyperconnected nodes with up to 3,473 and 4,141 connections in the gene-centric and proteoform-centric networks, respectively, while the most connected genes and proteoforms present 1,290 and 1,520 connections, respectively. Restricting small molecules to reaction-specific relationships allows considering the local function of small molecules without creating such hyperconnected nodes: the average degree of nodes is increased to 99 and to 109 for the gene- and the proteoform-centric networks, respectively, while the maximal degree increases to 2,361 and 2,376, and the maximal degree of small molecules remains 304 in both networks.

### PROTEOFORMS MODIFY THE LAYOUT OF CONNECTED COMPONENTS

*Connected components* are the maximal subnetworks in which all nodes of the component can reach each other through a path (**Figure 3**). The *Largest Connected Component (LCC)* of a network is the component with the highest number of nodes. In our analysis of Reactome, gene- and proteoform-centric networks showed similar relative size of the LCC (**Supplementary Table 6**). Given the hypothesis that nodes involved in the same biological function are connected in the pathway network, one expects that they should belong to the same connected component. Connected components can further be separated into subnetwork modules based on the topology of the network or based on their association with specific functions or diseases. Functional studies comparing such modules study the overlap between diseases to identify common molecular mechanisms or drug targets, and transfer knowledge of one module to the other. Thereby, nodes shared between biological processes have been suggested to be of particular interest for the study of disease mechanisms and treatment^2^.

**Figure 3.**
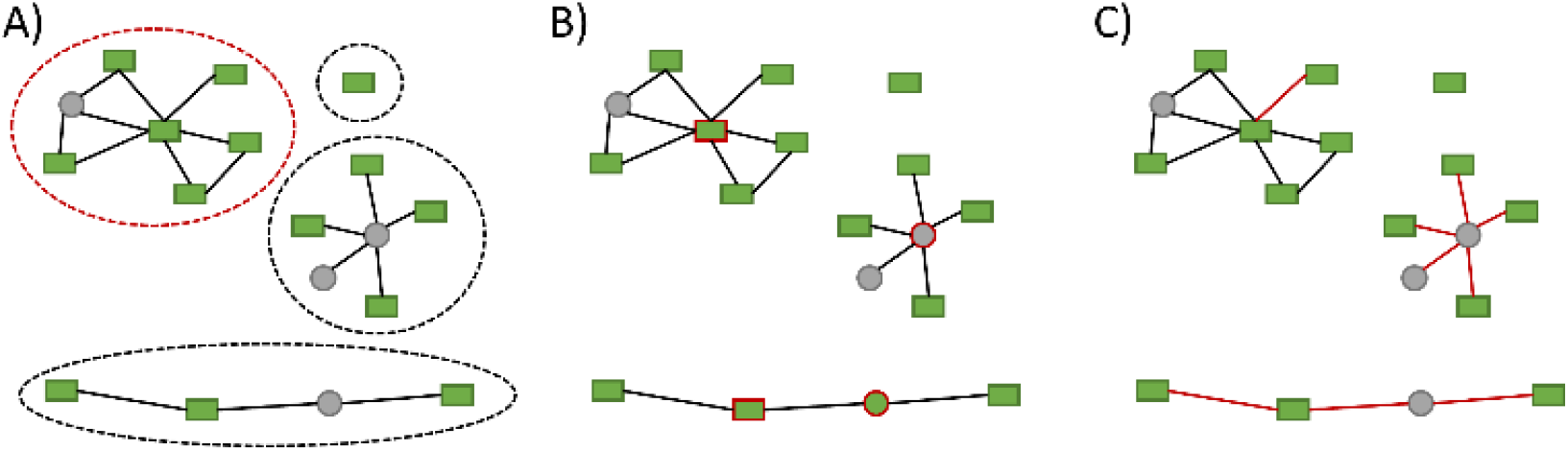
Illustration of graph theory concepts using hypothetical networks with proteoform nodes (green rectangles) and small molecule nodes (gray circles). A) *Connected components* of the network, each one surrounded with a dotted line. Largest connected component highlighted with red dotted line. B) *Articulation points*, nodes highlighted with a red border. C) *Bridges*, connections highlighted with red lines.

The proteoform interactome extends gene nodes into multiple proteoform nodes. Proteoforms resulting from variation of a single gene, called a proteoform family, may participate in disjoint sets of reactions in the network. If gene nodes are represented by multiple proteoforms participating in separate reactions or pathways, the overlap will only be observable at the gene level and not at the proteoform level. In other words, proteoforms from a single gene may be split over different modules and even different connected components. In this case, modules would intersect in the gene-centric representation of the network, but not in the proteoform-centric representation, where the different modules would be disconnected.

We found 497 proteins where at least one proteoform of the family participates in a biochemical reaction where the other members of the family are not involved. Identifying such a proteoform in a sample therefore provides pathway-specific information that is lost in a gene-centric representation, as in that case all reactions and pathways where any of the family members participate become indistinguishable. As an example, the human protein Peroxiredoxin-5 (P30044) has isoforms P30044-1 located at the Mitochondrial Matrix, and P30044-2 in the Cytosol. They differ in sequence, the second one missing the first 52 amino acids, and participate in separate reactions in different subcellular locations: “*PRDX5 reduces peroxynitrite to nitrite using TXN2*” and “*PRDX1,2,5 catalyzes TXN reduced + H2O2 => TXN oxidized + 2H2O”* respectively. In this case, a proteoform-centric module representation would distinguish the mitochondrial from the cytosol reaction, connecting them through the translocation and processing of P30044 into P30044-1, while a gene-centric representation would make both reactions indistinguishable.

### SMALL MOLECULES REDUCE THE PREVALENCE OF ISOLATED COMPONENTS AND NODES

Adding nodes representing small molecules considerably increases the percentage of nodes part of the LCC, from 85% to 98% in proteoform interactomes. Conversely, adding reaction-unique nodes for small molecules, rather than once for the whole interactome, prevents merging connected components when small molecules are the only nodes shared between reactions. By design, the number of connected components using reaction-unique small molecules is then greater than or equal to the number of connected components obtained when using small molecules, as displayed in Table 6, and consequently the LCCs are smaller. At the other end of the scale, some pathways contain proteins performing multiple roles in a reaction but not connected to other proteins, leading to isolated nodes only connected to themselves in the network. This may happen when different isoforms or proteoforms of the same protein participate in the reaction with different roles, resulting in the gene centric representation being a single node interacting with itself while the proteoform-centric representation would show a module composed of multiple nodes. We found 1,665 and 1,696 isolated nodes for the gene- and proteoform-centric networks, respectively, showing an overall stable number of isolated nodes. For example, the reactions sustaining Vitamin B1 (thiamin) metabolism (**Figure 4**) yield isolated nodes that stay isolated even in the proteoform-centric representation.

**Figure 4.**
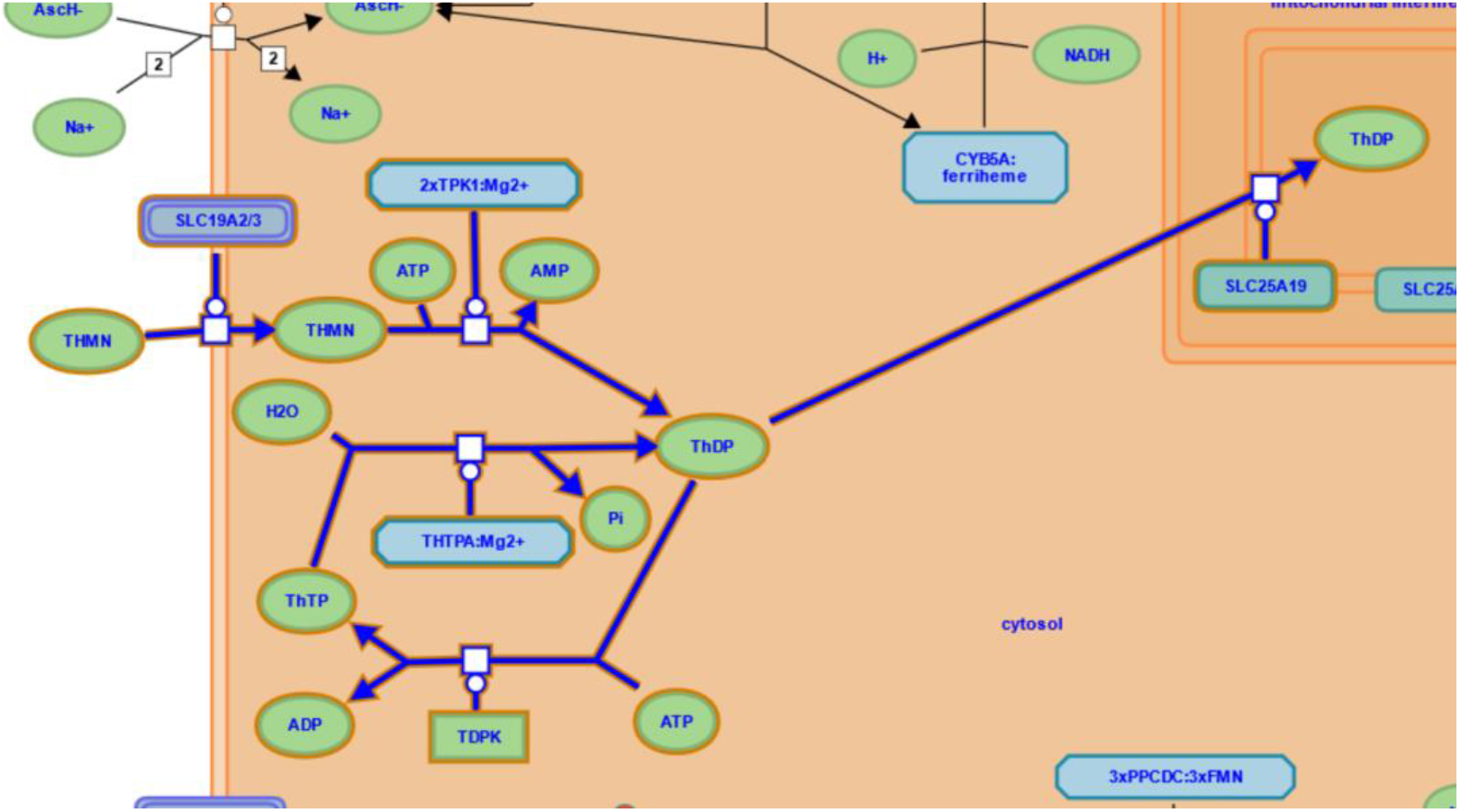
Section of Reactome Pathway diagram of “*Vitamin B1 (thiamin) metabolism*” (R-HSA-196819). White squares represent reactions composing the pathway. Green ovals are small molecules. Green rectangles represent protein molecules. Blue rectangles with chopped corners are complex molecules. Dark blue arrows show relationship between reactants (inputs) and products (outputs) of the reactions following the direction of the arrow from input to output. Molecules connected with a white circle on the arrow represent catalyst molecules.

Adding small molecules reduces the number of isolated nodes to 164 and 171, respectively. Among the 10,976 accessioned entity nodes, 2,789 (25 %) are connected only through small molecules. When considering 1,119 pathways, 226 displayed less isolated nodes when considering small molecules. Conversely, for 39 pathways there were more isolated nodes when adding small molecules. When the studied network is sparse or with many disconnected nodes, it thus becomes useful to include small molecules. They show indirect ways to reach one gene from another through reactions, yet the relevance of connecting two distant proteins by a small molecule can be questioned. Reaction-unique small molecules provide a balance between reducing the number of isolated nodes while not connecting nodes across different pathways. They allow connecting otherwise isolated nodes through a path that alternates between accessioned entities and reaction-relevant small molecules, while preserving the disconnection of pathways and components.

### ARTICULATION POINTS AND BRIDGES

Articulation points and bridges are respectively nodes and connections that, if removed, break one connected component into two or more components (**Figure 3**). They are thus important members of the network, maintaining the connection between otherwise disconnected clusters of nodes. The higher the prevalence of articulation points and bridges, the less robust the network. We therefore investigated whether the prevalence of bridges and articulation points changed from a gene-centric to a proteoform-centric representation. However, as detailed in **Supplementary Table 8** and **Supplementary Table 9** for articulation points and bridges, respectively, adding proteoform annotation does not substantially change the share of articulation points (from 2.43 % to 2.46 % of nodes). Articulation points in the gene-centric network either stay articulation points in the proteoform-centric network or become more connected due to the multiplicity of proteoform nodes in a proteoform family. Therefore, proteoforms do not yield to more isolated nodes but may create more connected components (**Supplementary Table 6**). This indicates that, although proteoform annotation increases the connectivity in the network, it is mainly through within-component connectivity.

Given the ubiquitous nature of some small molecules, which participate in many pathways across many contexts, and the increased connectivity that they induce, as observed in the previous sections, it can be anticipated that they create new connections between connected components. Indeed, adding small molecules reduced the prevalence of bridges and articulation points. In proteoform-centric networks they reduce from 351 (2,46 %) to 254 (1,56 %). In the network extended with small molecules, 40 % of articulation points were small molecules and 60 % accessioned entities (**Supplementary Table 8**). Conversely, when restricting the role of small molecules to single reactions, the number of bridges was tripled. Thus, adding reaction-specific single molecules improved the connectivity of the network through single reactions, that are biologically more specific, but less robust. Small molecules do not perform biological processes on their own, they need to interact with accessioned entities. Therefore, they are rarely the only shared node between steps of pathways, resulting in being articulation points less frequently. Adding reaction-specific small molecules also has the effect of increasing the percentage of proteoforms that are articulation points and increasing the percentage of bridges going out of small molecule nodes, from 3.55 % to 10.70 % (**Supplementary Table 9**).

We investigated the changes in prevalence and nature of bridges and articulation points at the level of pathways. **Supplementary Table 10** and **Supplementary Table 11** detail the averaged values among all pathways in Reactome considered individually. The share of articulation points considering pathways one by one is slightly higher than when considering the complete interactome, highlighting how interactomes aggregate pathways, overlapping the connections of nodes in different contexts. Once again, accessioned entities are more often articulation points than small molecules, demonstrating their key role in biological processes. Nevertheless, small molecules still represent one third of articulation points. Even per pathway, the tendency of small molecules to reduce the percentage of bridge connections is clear, confirming the important role of small molecules for the connectivity of the network at both local and global levels.

Bridges connect more than twice as often accessioned entities rather than small molecules, and the prevalence increases when studying per pathway than for the complete interactome. This increase can be interpreted as the connections of proteoforms conveying more unique information, whereas small molecules may connect more diverse types of other molecules. Reaction-unique small molecules are expected to be articulation points more frequently than regular small molecules, but no difference was found on average. Reaction-unique small molecules increase the total number of articulation points by increasing the percentage of accessioned entity nodes that are articulation points. This is due to the smaller average node degree of the reaction-unique small molecules, compared to the regular small molecules. Hence, when they connect to an accessioned entity, they may convert that accessioned entity into an articulation point.

## CONCLUSIONS

Interaction networks are a useful representation of biological processes, *e.g*., to study if biological entities are functionally related. We demonstrate multiple ways to build interaction networks from the Reactome pathway knowledgebase, resulting in networks with very different general and local properties. Such differences in network topology are likely to influence the biological interpretation of experimental data. In particular, we explored the possibility of adding proteoforms and small molecules, which are usually not considered when building interaction networks, despite playing essential roles in biological processes.

We show that extending the representation of proteins using isoforms and post-translational modifications has an impact on the structure of biological networks, but that this information is only available for a subset of the proteins. This results in highly connected interfaces between single proteins with no proteoform annotation and proteoforms from the same family. We also demonstrate how small molecules such as metabolites alter the structure of the networks. Including them can help connecting sparse areas of the network, but it can also result in highly connected nodes with little to no biological relevance. As a compromise, we propose to take advantage of the annotation of pathway databases to restrict the interactions of such molecules.

With this study we further compared the results of topological analyses conducted to the level of the entire network and when considering one pathway at a time. Each approach yielded different results, showing how local and global properties of the network differ. This highlighted how the arbitrary representation of pathways may alter the perception of the connectedness of biological entities, hiding inter-pathway connections. Overall, our results point towards the importance of using the rich information contained in pathway databases to contextualize network analyses while also highlighting the difficulty to provide an unbiased representation of interaction networks, both locally and globally.

## DISCUSSION

This study investigates the impact of changing the network representation through the inclusion of proteoforms and small molecules. We base these findings on the Reactome knowledgebase, which contains rich information on biological pathways. Due to the high level of detail on biochemical reactions required to build such networks, functional annotation on proteoforms and small molecules is still scarce. The rapid pace in increase of functional knowledge indicates that such analyses will become increasingly powerful. As the interactome becomes more connected, refining its representation using the rich information available in pathway knowledgebases represents a promising avenue to tease apart densely connected functional regions.

Factors such as analytical challenges, research interest, and literature curation lead to some proteins or pathways to be better annotated than others. The better annotated pathways give a more detailed representation of the biological processes, while understudied proteins or pathways have a much less mature representation or even remain undiscovered. Such biases have a strong influence on the representation of the biological processes involved, and dramatically alter the ability to conduct refined studies such as proteoform-level network analyses. The disparity in biological functional knowledge is a strong limitation of the field, yielding to a network where some processes yield densely connected subnetworks of proteoforms and small molecules, while others are only represented by sparse disconnected gene names – when any information is available at all.

Technologies to identify proteoforms and small molecules are improving constantly but integrating these biological entities in pathways at scale poses numerous challenges. It is therefore important to develop new biological network analysis approaches that can handle the heterogeneity in pathway annotation without losing the rich information gathered by the scientific community. One can envision that such approaches will be generalizable to hybrid networks combining pathway knowledgebases with interaction networks derived from experiments or text mining.

Constructing an interaction network using refined information like proteoforms or small molecules is even more challenging using multiple sources of data. Functional annotations often refer only to gene or protein accessions^19^, hence overlooking post-translational regulatory mechanisms central to many biological processes. The broad adoption of proteoforms in the representation of biological processes is essential to generalize the approaches presented in this study, and hence allow the refinement of the representation of biological processes, which will eventually provide biomedical researchers with more powerful tools to interpret their data.

## Acknowledgments

This work was supported by the Research Council of Norway (project #301178 to M.V.) and by the Bergen Research Foundation (project #BFS2016REK02 to H.B.B. and H.B.).

## Competing interests

The authors declare no competing interests.

## METHODS

Reference knowledge to conduct the analysis was obtained from the Reactome graph database (version 80). The database dump file (reactome.org/download-data) was loaded and run using Neo4j Desktop 1.4.2 to Neo4j Graph Database Manager 4.4.5. The analysis scripts were implemented using Python 3.10.2 organized as Jupyter notebooks. They communicated with the database management system using the Neo4j Python Driver Manual version 4.4.2.

All the code used to construct the networks and replicate the topological analysis is publicly available at the public repository: github.com/PathwayAnalysisPlatform/ProteoformNetworks

The Reactome graph database data model is organized as nodes and relationships with properties and labels. The main nodes we used were *Event* nodes, which involve the transformation of input nodes to output nodes in one or multiple steps. We queried for two types of event nodes: *Pathway* and *ReactionLikeEvents*. *ReactionLikeEvents* convert input entities to output entities in one step, while *Pathways* group sets of *ReactionsLikeEvents*. Each event has participant molecules which perform roles of input (reactant), output (product), regulators and catalyzers (enzymes). The data model represents events occurring in sequence by annotating the output of the first event as input of the second event.

Participants of reactions are physical entities, which typically are of two types: accessioned sequences entities (genes, transcripts, or proteins) or small molecules (metabolites, water, etc.). Accessioned sequences stand for those molecules which have a standard identifier for each sequence pattern, typically nucleotide-based sequences (DNA pieces like genes) or amino acid-based sequences (proteins). Genes are annotated with HUGO gene nomenclature identifiers^16^, while proteins have UniProt^20^ accession numbers. Small molecules also have unique identifiers from the Chemical Entities of Biological Interest (ChEBI) database^21^. These refer to the chemical element or compound rather than a sequence molecule.

When this information is available, accessioned sequence participants are annotated with their isoform and the minimal set of post-translational modifications necessary to perform their role in the biological event. Combining the set of modifications and isoform sequence, we built a theoretical proteoform state in which the gene products need to be present to participate in a given reaction. Participants of events may also be entity sets or complexes. Entity sets stand for groups of entities which may be used almost interchangeably with the desired role in the biological event. For example, multiple proteins may interchangeably perform the same role in a reaction, *e.g*., catalyzing a reaction. Complexes are the conjunction of multiple molecules into a single unit. The members of the complex may be of all other types of participants, *i.e*., accessioned sequences, proteins, metabolites or even complexes.

We constructed gene- and proteoform-centric interaction network representations of the Pathways in Reactome by taking participating entities of reactions as nodes of the network, as in previous studies^3,15^. For the gene-centric representation, all physical entities associated with a given gene and each of its associated UniProt protein accessions are represented by a single node, *i.e*., merging all protein products, isoforms and proteoforms into one node. For the proteoform-centric representation, we represent each proteoform with a separate node. We take the associated protein accession, the isoform, and set of post-translational modifications annotated to represent a single proteoform. Then, all physical entities yielding the same isoform with the same sequence modification combinations are represented by a single node. For both the gene- and proteoform-centric networks we constructed two alternative networks which additionally considered the small molecule participants of reactions; the first alternative adds a single node for each small molecule, the second alternative adds a node for each small molecule to every reaction in which the given small molecule participates.

Once nodes are defined, we set a connection between two nodes when they perform a role in the same reaction, such as input and output. To construct a complete interactome we process all pathways with all their respective reactions to obtain their nodes and connections. We do not repeat nodes, but instead aggregate their connections obtained from each pathway. The resulting network contains all annotated genes or proteoforms for humans in the pathway database.

Networks were represented using the Networkx library version 2.7.1 for Python. The library allowed the calculation of size, articulation points, bridges, and connected components. The robustness of the network was calculated through percolation analysis resulting in a percolation curve, which shows the average size of the LCC, called *giant component*, when random nodes or connections are removed.

There is usually a point when the size of the LCC collapses rapidly, that represents the percolation threshold, indicating the average size of modules that can be observed in the network^2^.

---

**Supplementary Table 1.**
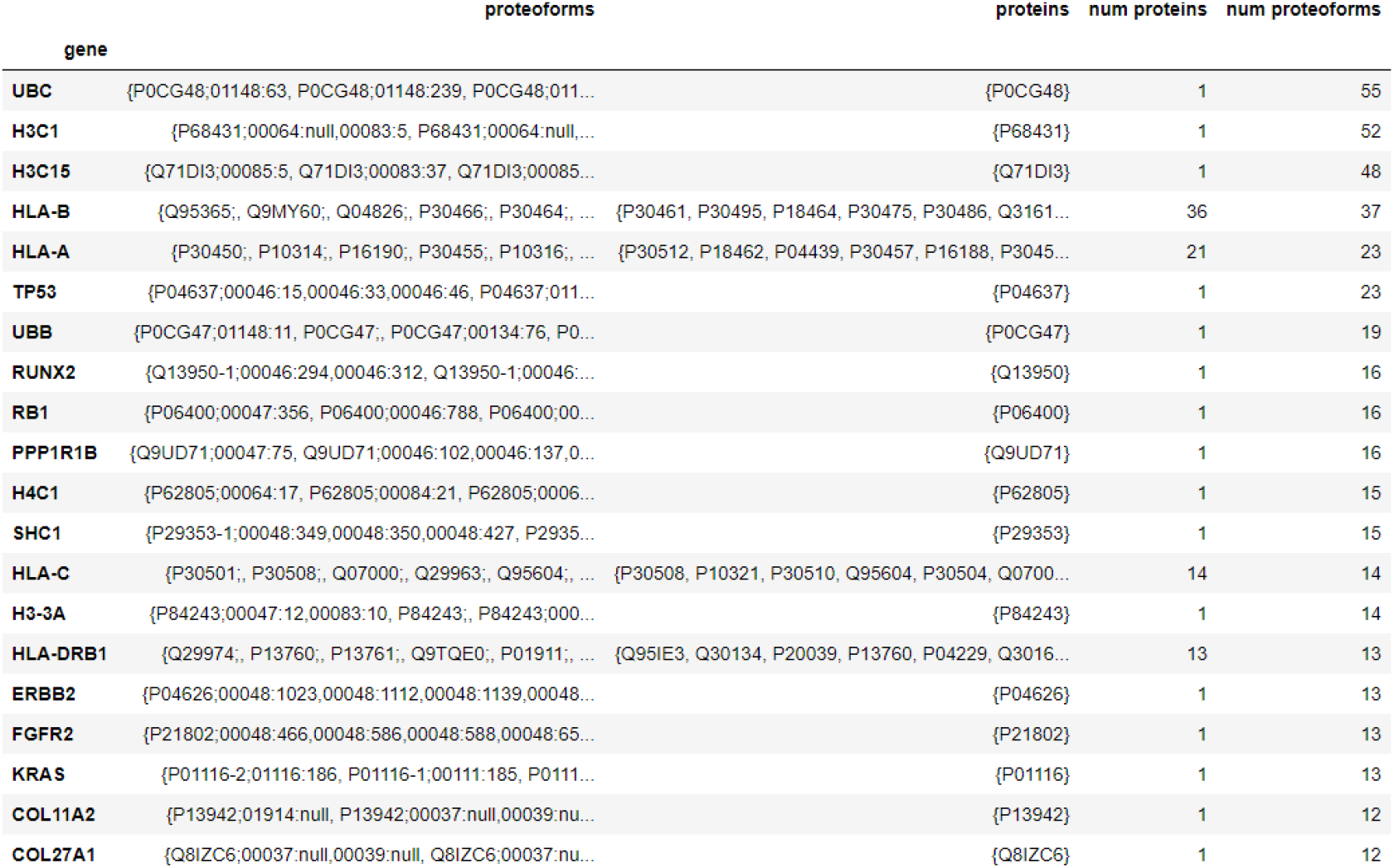
Top 20 genes with most proteoforms products participating in Reactome pathways. For each gene the number of protein and proteoform products is shown, along with some protein and proteoform examples. Proteins are represented by their UniProt Accessions; Proteoforms by the protein isoform variant and a set of post translational modifications; and each modification as a PSIMOD identifier paired with an integer coordinate indicating its location on the protein sequence. If no localization information is known, “null” replaces the localization.

**Supplementary Table 2.** Sizes of six alternative interactome networks resulting from combining entity level (genes, proteoforms) and three options to consider small molecule nodes. Sizes are shown as number of connections (interactions) and number of nodes.

|  |  | Interactions | Nodes |
| --- | --- | --- | --- |
| Small Molecules | Entity Level |  |  |
| Not Included | genes | 366208 | 10976 |
|  | proteofoms | 590415 | 14246 |
| Included | genes | 451490 | 13033 |
|  | proteofoms | 681891 | 16303 |
| Reaction-Unique Included | genes | 808212 | 40575 |
|  | proteofoms | 1047542 | 43845 |

**Supplementary Table 3.** Descriptive summary statistics on the node degree for the different interactome networks resulting from combining entity level (genes, proteoforms) and three options to consider small molecule nodes. Degree values are shown in separate columns for the two types of nodes: accessioned entities (AE) and small molecules (SM). Columns show quartiles of node degree values separating the lowest 25% as Q1, median at 50% as Q2, value setting the top 75% as Q3 and the maximum value as Q4.

|  |  | Q1 AE | Q2 AE | Q3 AE | Q4 AE | Avg AE | Q1 SM | Q2 SM | Q3 SM | Q4 SM | Avg SM |
| --- | --- | --- | --- | --- | --- | --- | --- | --- | --- | --- | --- |
| Small Molecules | Entity Level |  |  |  |  |  |  |  |  |  |  |
| Not Included | genes | 3.00 | 22.00 | 91.00 | 1,243.00 | 66.73 | 0.00 | 0.00 | 0.00 | 0.00 | 0.00 |
|  | proteofoms | 4.00 | 23.00 | 109.00 | 1,474.00 | 82.89 | 0.00 | 0.00 | 0.00 | 0.00 | 0.00 |
| Included | genes | 5.00 | 25.00 | 98.00 | 1,290.00 | 72.88 | 4.00 | 7.00 | 20.00 | 3,473.00 | 50.09 |
|  | proteofoms | 6.00 | 26.00 | 116.00 | 1,520.00 | 88.06 | 4.00 | 7.00 | 20.00 | 4,141.00 | 53.10 |
| Reaction-Unique Included | genes | 5.00 | 30.00 | 115.00 | 2,361.00 | 99.23 | 2.00 | 4.00 | 10.00 | 304.00 | 17.81 |
|  | proteofoms | 6.00 | 30.00 | 128.00 | 2,376.00 | 108.99 | 2.00 | 4.00 | 10.00 | 304.00 | 18.32 |

**Supplementary Table 4.** Degree comparison of nodes from the gene and proteoform interactomes. Each row shows a proteoform with its source gene, both with their respective degree.

|  | Proteoform Degree | Gene | Gene Degree |
| --- | --- | --- | --- |
| Proteoform |  |  |  |
| Q9UBT2;00211:173 | 9 | UBA2 | 6 |
| P68431;00064:null,00083:28 | 10 | H3C1 | 348 |
| Q17RW2;00037:null,00038:null,00039:null | 544 | COL24A1 | 66 |
| Q14994-1; | 85 | NR1I3 | 52 |
| P39059;00037:null,00039:null | 536 | COL15A1 | 67 |
| Q7L9L4;00047:12,00047:35 | 13 | MOB1B | 8 |
| P60484;01148:13,01148:289 | 7 | PTEN | 34 |
| P56524;01149:559 | 39 | HDAC4 | 58 |
| Q14155-1; | 2 | ARHGEF7 | 142 |
| P24941;00048:15 | 5 | CDK2 | 158 |
| P50542-1; | 65 | PEX5 | 67 |
| P22607;00048:577,00048:647,00048:648,00048:724,00048:760,00048:770 | 30 | FGFR3 | 91 |
| Q92614;00048:599,00048:768,00048:955,00048:969 | 27 | MYO18A | 37 |
| P05161;00134:157 | 25 | ISG15 | 60 |
| P36957;00127:110 | 14 | DLST | 13 |
| Q96J94;00076:14,00078:49,00078:53,00078:370 | 3 | PIWIL1 | 3 |
| P11216;00128:681 | 91 | PYGB | 92 |
| P19419;00046:324,00046:383,00046:389,00046:422,00047:336 | 8 | ELK1 | 7 |
| Q9BXL7;00046:552,00046:645 | 2 | CARD11 | 14 |
| Q07092;00038:null,00039:null,01914:null | 538 | COL16A1 | 63 |

**Supplementary Table 5.**
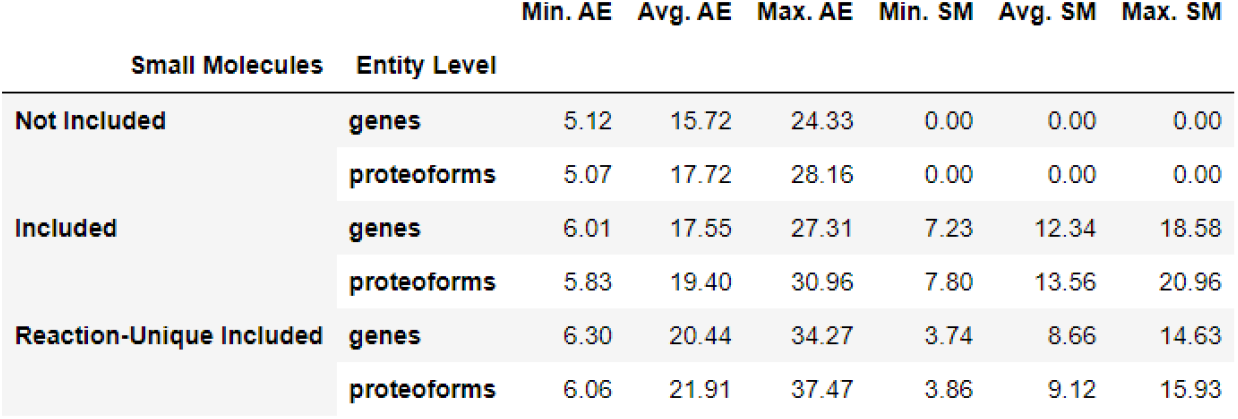
Descriptive summary statistics on the node degree per pathway. Values for each pathway are taken from six networks resulting from combining entity level (genes, proteoforms) and three options to consider small molecule nodes. Degree values are shown in separate columns for the two types of nodes: accessioned entities (*AE*) and small molecules (*SM*). Values refer only to pathways where proteoforms annotated with isoform or post-translational modifications participate.

|  |  | Min. AE | Avg. AE | Max. AE | Min. SM | Avg. SM | Max. SM |
| --- | --- | --- | --- | --- | --- | --- | --- |
| Small Molecules | Entity Level |  |  |  |  |  |  |
| Not Included | genes | 5.12 | 15.72 | 24.33 | 0.00 | 0.00 | 0.00 |
|  | proteoforms | 5.07 | 17.72 | 28.16 | 0.00 | 0.00 | 0.00 |
| Included | genes | 6.01 | 17.55 | 27.31 | 7.23 | 12.34 | 18.58 |
|  | proteoforms | 5.83 | 19.40 | 30.96 | 7.80 | 13.56 | 20.96 |
| Reaction-Unique Included | genes | 6.30 | 20.44 | 34.27 | 3.74 | 8.66 | 14.63 |
|  | proteoforms | 6.06 | 21.91 | 37.47 | 3.86 | 9.12 | 15.93 |

**Supplementary Table 6.** Descriptive summary statistics on the connected components (CCs) for the interactome networks resulting from combining entity level (genes, proteoforms) and three options to consider small molecule nodes. Largest Connected Component (LCC) and Smallest Connected Component (SCC) sizes are evaluated using the number of nodes. Relative size of the LCC represents the fraction of nodes in the complete network that are also members of the LCC.

|  |  | Num. CCs | Size of LCC | Relative size of LCC | Average size of CCs | Size of SCC | Num. isolated nodes |
| --- | --- | --- | --- | --- | --- | --- | --- |
| Small Molecules | Entity Level |  |  |  |  |  |  |
| Not Included | genes | 1774 | 8967 | 0.82 | 6.19 | 1 | 1665 |
|  | proteoforms | 1819 | 12091 | 0.85 | 7.83 | 1 | 1696 |
| Included | genes | 194 | 12792 | 0.98 | 67.18 | 1 | 164 |
|  | proteoforms | 202 | 16053 | 0.98 | 80.71 | 1 | 171 |
| Reaction-Unique Included | genes | 2445 | 30153 | 0.74 | 16.60 | 1 | 1160 |
|  | proteoforms | 2480 | 33107 | 0.76 | 17.68 | 1 | 1167 |

**Supplementary Table 7.**
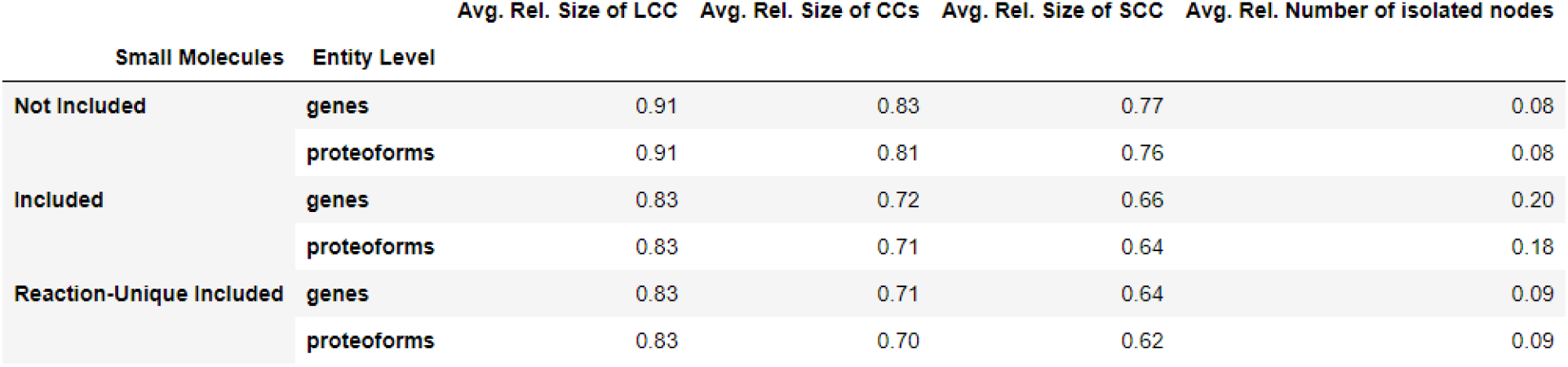
Descriptive summary statistics on connected components (CCs) per pathway. Values for each pathway are taken from six alternative networks resulting from combining entity level (genes, proteoforms) and three options to consider small molecule nodes. Relative size refers to the fraction of nodes in the complete network that are also members of a connected component. Values are an average of the values per pathway, considering only pathways where proteoforms annotated with isoform or post-translational modifications participate.

**Supplementary Table 8.** Descriptive summary statistics on the prevalence of articulation points for the networks resulting from combining entity level (genes, proteoforms) and three options to consider small molecule nodes. Columns show the total number of articulation points in each network, the percentage of nodes in the network that are articulation points, then by node type: accessioned entities (AE) and small molecules (SM).

|  |  | Art. Points | % Art. Points | AE | % AE | SM | % SM |
| --- | --- | --- | --- | --- | --- | --- | --- |
| Small Molecules | Entity Level |  |  |  |  |  |  |
| Not Included | genes | 267 | 2.43 | 267 | 2.43 | 0 | 0.00 |
|  | proteoforms | 351 | 2.46 | 351 | 2.46 | 0 | 0.00 |
| Included | genes | 244 | 1.87 | 143 | 1.10 | 101 | 0.77 |
|  | proteoforms | 254 | 1.56 | 151 | 0.93 | 103 | 0.63 |
| Reaction-Unique Included | genes | 2012 | 4.96 | 1600 | 3.94 | 412 | 1.02 |
|  | proteoforms | 2050 | 4.68 | 1638 | 3.74 | 412 | 0.94 |

**Supplementary Table 9.** Descriptive summary statistics on the prevalence of bridges for the networks resulting from combining entity level (genes, proteoforms) and three options to consider small molecule nodes. Columns show the total number of bridges, the percentage of connections in each network that are bridges, the percentage of connections of a node that are bridges, then by node type: accessioned entities (*AE*) and small molecules (*SM*).

|  |  | Bridges | % Bridges | % Bridges out of AE | % Bridges out of SM |
| --- | --- | --- | --- | --- | --- |
| Small Molecules | Entity Level |  |  |  |  |
| Not Included | genes | 511 | 0.14 | 5.41 | 0.00 |
|  | proteforms | 582 | 0.10 | 4.81 | 0.00 |
| Included | genes | 523 | 0.12 | 4.49 | 3.53 |
|  | proteforms | 527 | 0.08 | 3.52 | 3.55 |
| Reaction-Unique Included | genes | 3444 | 0.43 | 5.87 | 11.07 |
|  | proteforms | 3356 | 0.32 | 4.63 | 10.79 |

**Supplementary Table 10.** Descriptive summary statistics on the prevalence of articulation points per pathway. Values for each pathway are taken from six networks resulting from combining entity level (genes, proteoforms) and three options to consider small molecule nodes. Values are averaged per pathway, considering only pathways where proteoforms annotated with isoform or post-translational modifications participate. Columns show the total number of articulation points in each network, the percentage of nodes in the network that are articulation points, then by node type: accessioned entities (*AE*) and small molecules (*SM*).

|  |  | Art. Points | % Art. Points | AE | % AE | SM | % SM |
| --- | --- | --- | --- | --- | --- | --- | --- |
| Small Molecules | Entity Level |  |  |  |  |  |  |
| Not Included | genes | 0.74 | 3.03 | 0.74 | 3.03 | 0.00 | 0.00 |
|  | proteforms | 1.02 | 3.92 | 1.02 | 3.92 | 0.00 | 0.00 |
| Included | genes | 0.86 | 2.86 | 0.58 | 2.05 | 0.28 | 0.80 |
|  | proteforms | 0.91 | 2.65 | 0.62 | 1.88 | 0.29 | 0.78 |
| Reaction-Unique Included | genes | 1.67 | 3.68 | 1.45 | 3.31 | 0.23 | 0.37 |
|  | proteforms | 1.71 | 3.42 | 1.49 | 3.06 | 0.23 | 0.36 |

**Supplementary Table 11.** Descriptive summary statistics on the prevalence of bridges per pathway. Values of each pathway are taken from six networks resulting from combining entity level (genes, proteoforms) and three options to consider small molecule nodes. Values are averaged per pathway, considering only pathways where proteoforms annotated with isoform or post-translational modifications participate. Columns show the total number of bridges, the percentage of connections in each network that are bridges, the percentage of connections of a node that are bridges, then by node type: accessioned entities (*AE*) and small molecules (*SM*).

|  |  | Bridges | % Bridges | % Bridges out of AE | % Bridges out of SM |
| --- | --- | --- | --- | --- | --- |
| Small Molecules | Entity Level |  |  |  |  |
| Not Included | genes | 2.03 | 0.10 | 10.94 | 0.00 |
|  | proteforms | 2.28 | 0.10 | 11.14 | 0.00 |
| Included | genes | 1.43 | 0.04 | 6.66 | 2.74 |
|  | proteforms | 1.42 | 0.03 | 5.43 | 2.38 |
| Reaction-Unique Included | genes | 3.17 | 0.05 | 7.72 | 7.45 |
|  | proteforms | 3.08 | 0.04 | 6.37 | 6.91 |

## Notes

### Competing Interest Statement

The authors have declared no competing interest.

